# *TP53* somatic mutations in Asian breast cancer are associated with subtype-specific effects

**DOI:** 10.1101/2022.03.31.486643

**Authors:** Mohana Eswari Ragu, Joanna Mei Ch’Wan Lim, Pei-Sze Ng, Cheng-Har Yip, Pathmanathan Rajadurai, Soo-Hwang Teo, Jia-Wern Pan

## Abstract

Recent genomics studies of breast cancer in Asian cohorts have found a higher prevalence of *TP53* mutations in Asian breast cancer patients relative to Caucasian patients. However, the effect of *TP53* mutations on Asian breast tumours has not been comprehensively studied. Here, we report an analysis of 492 breast cancer samples from the Malaysian Breast Cancer (MyBrCa) cohort where we examined the impact of *TP53* somatic mutations in relation to PAM50 subtypes by comparing whole exome and transcriptome data from tumours with mutant and wild type *TP53*. We found that the magnitude of impact of *TP53* somatic mutations appears to vary between different subtypes. *TP53* somatic mutations were associated with larger and more consistent differences in HR deficiency scores as well as transcriptional alterations in the luminal A and luminal B subtypes compared to the basal-like and Her2-enriched subtypes. The only pathways that were consistently dysregulated when comparing tumours with mutant and wild type *TP53* across the different subtypes were the mTORC1 signaling and glycolysis pathways. These results suggest that therapies that target *TP53* or other downstream pathways may be more effective against luminal A and B tumours in the Asian population.

## Introduction

Breast cancer continues to be the most common cancer in Asian women, and the incidence rates of breast cancer among Asian women is predicted to increase in the coming years (Sung et al., 2021). Breast cancer in Asian women appears to have a distinct clinical presentation, with a younger age of onset and increased frequencies of HER2+ tumors relative to European populations (Pan et al., 2020). Increasingly, genomics studies of Asian breast tumours suggest that they have a distinct molecular profile as well, with a more active immune microenvironment and higher frequencies of *TP53* somatic mutations (Kan et al., 2018; Pan et al., 2020; Yap et al., 2018; Zhu et al., 2019). Interestingly, even though *TP53* somatic mutations are generally more common in ER-tumours, the increased prevalence of *TP53* somatic mutations in Asian women relative to Caucasian women appears to be more pronounced in ER+ tumour (Pan et al. 2020).

The p53 protein encoded by *TP53* is involved in wide range of cellular stress responses, with a wide range of downstream effects including cell cycle arrest, apoptosis, senescence, DNA repair or changes in metabolism (Liu et al., 2013). The significance of TP53 in tumor development is demonstrated that TP53 mutations befall in almost all of human cancers and these mutations are typically missense and are largely situated in exons 5–8, covering the DNA-binding domain of the protein (Petitjean et al., 2007). The TP53 mutations possibly instigate whole or fractional loss of protein function or gain of function (GOF) (Muller and Vousden,2013; Oren and Rotter, 2010).

The frequency and type of TP53 somatic mutations varies across the breast cancer PAM50 molecular subtypes, and is most common in the basal-like tumour subtypes and lowest in Luminal A subtype (Silwal-Pandit et al., 2017). Early studies in Caucasian populations suggested that the prognostic significance of *TP53* mutations is independent of hormone receptor status (Olivier et al., 2006); however, more recent studies suggest that *TP53* somatic mutations may have subtype-specific impacts on prognosis (Silwal-Pandit et al., 2014). Specifically, mutations in *TP53* were associated with increased mortality in patients with luminal B, HER2-enriched, and normal-like tumors but not in patients with luminal A and basal-like tumors (Silwal-Pandit et al., 2014). In contrast, *TP53* somatic mutations in Asian breast cancer patients have been associated with slightly better overall survival in ER+ patients, but not in ER-patients (Pan et al., 2020). Thus, further exploration of the effect of *TP53* somatic mutation in each PAM50 subtype across different populations may uncover population-specific data that could clarify the role of *TP53* as a predictive or prognostic biomarker for breast cancer.

Here we report an analysis of *TP53* somatic mutations in a cohort of 489 Malaysian female breast cancer patients of all ages. In this analysis, we compare various molecular characteristics between tumour samples with mutant *TP53* and wild type *TP53* across the PAM50 molecular subtypes using whole exome and whole transcriptome data. We identify significant differences in HR deficiency scores, gene expression, and molecular pathways that vary between different breast cancer subtypes, but appear to be particularly pronounced in luminal A and luminal B tumours. These results may have clinical implications for the current and future use of therapies that target *TP53* or other downstream pathways in the Asian population.

## Results

### Cohort characteristics

The MyBrCa tumour cohort comprises of 560 Malaysian breast cancer patients recruited from single Malaysian private hospital (Subang Jaya Medical Centre). From this initial cohort, we excluded patient samples without RNAseq data as well as patient samples with *TP53* mutations classified as benign, likely benign, or uncertain significance, for a final dataset of 492 samples from 489 patients (3 cases of bilateral tumours). Genetically, this cohort overlaps with other East/Southeast Asian populations according to genotyping analysis (Ho et al., 2020). Of the 492 samples, 199 samples were classified as having mutant *TP53* (pathogenic/likely pathogenic somatic mutations), and the remaining 293 samples were classified as wild type *TP53* to serve as our control group.

Table 1 shows the relationship between *TP53* somatic mutations and patients’ clinical characteristics, after excluding patients with bilateral tumours. We found significant differences between patients with wild type vs mutant *TP53* with respect to PAM50 subtypes, tumor grade and histological subtype. Samples with mutant *TP53* were associated with basal-like, Her2-enriched tumours and higher tumour grade, and negatively associated with lobular carcinomas (Table 1). There were no significant differences observed between the two groups with respect to age at cancer diagnosis and tumour stage.

**Table 1.**
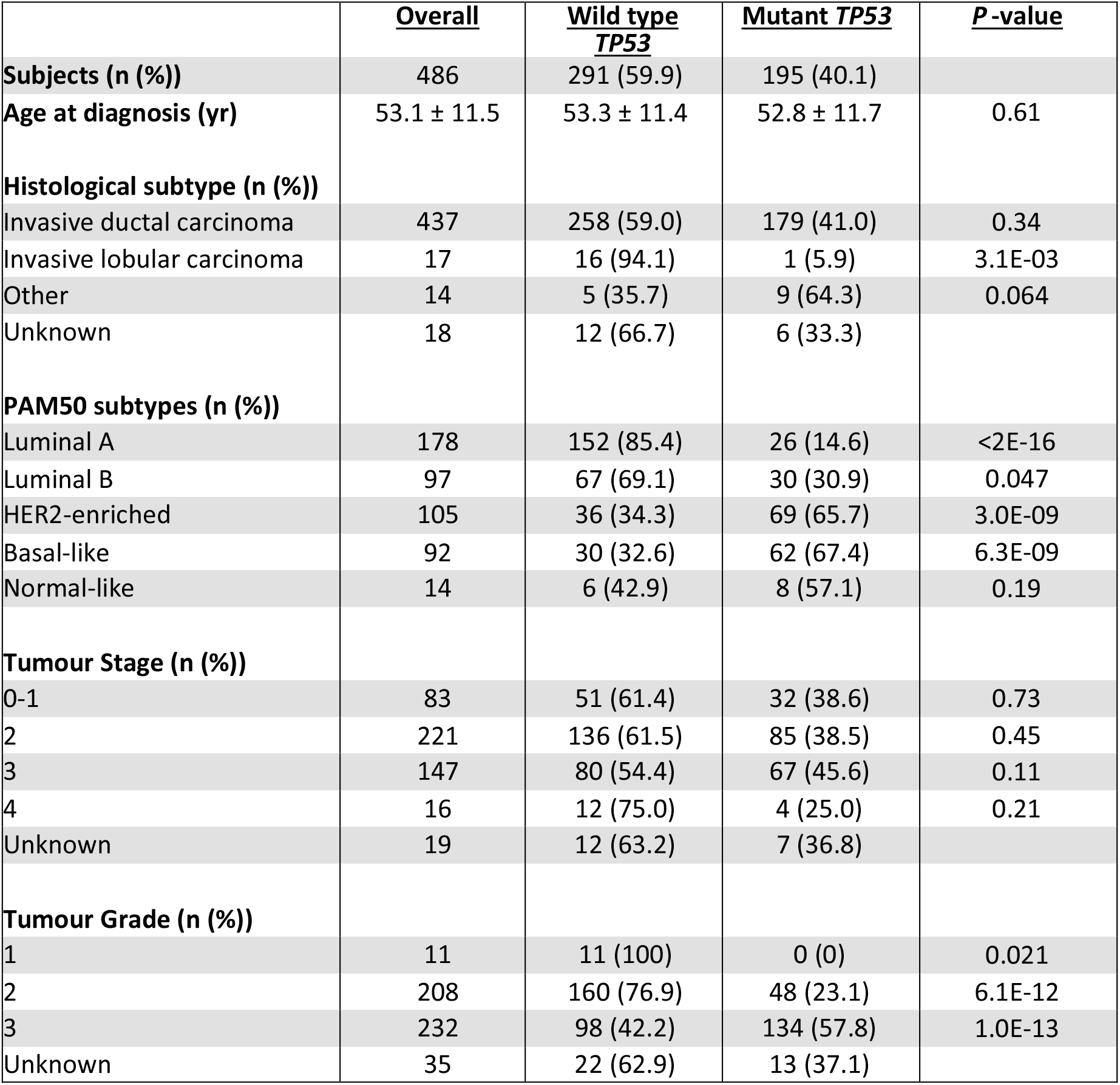
Cohort characteristics. Statistical significance determined with Student’s t-test or chi-square tests, excluding “Unknown” samples.

### Characterization of TP53 mutations

To investigate the role of *TP53* mutations in the MyBrCa cohort, we began our analysis by comparing the location and distribution of deleterious mutations in the *TP53* gene between the main PAM50 subtypes. The majority of the mutations were substitutions (missense and nonsense mutations (n = 139, 69.8%), followed by frame shift indels (n = 41, 20.6%) and splice site variants (n = 13, 6.5%). However, there was no significant difference in the location of the mutations, with 84.7% of mutations occurring in the DNA binding domain of *TP53* for luminal A and luminal B samples and 86.2% for basal-like and Her2-enriched samples (Supplementary Figure 1). The R273C and R175H mutations, known to be common hotspot mutations in *TP53*, can be observed across all the subtypes at similar frequencies (Supplementary Table 1).

### Mutational profiles of tumours with *TP53* somatic mutations

Next, we examined the mutational profile of tumour samples for differences associated with mutant *TP53* across subtypes, beginning with an analysis of tumour mutational burden. Using WES data, we established the total number of somatic mutations (small insertion–deletions (indels) and single nucleotide variants (SNVs)) for each tumour sample and additionally included tumours with known germline *BRCA* mutations as a positive control. As expected, tumours with germline *BRCA* mutations had a significantly higher number of somatic mutations compared to non-carriers (*p* = 0.001, Figure 1a). Similarly, tumours with *TP53* somatic mutations overall had a significantly higher numbers of somatic mutations compared to tumours with wild type *TP53* (*p* < 1e-5, Figure 1a). However, although the number of somatic mutations was numerically higher in mutant *TP53* samples compared to wild type *TP53* samples in each of the breast cancer subtypes, the results were statistically significant only luminal B samples (p=0.008, Figure 1a).

**Figure 1.**
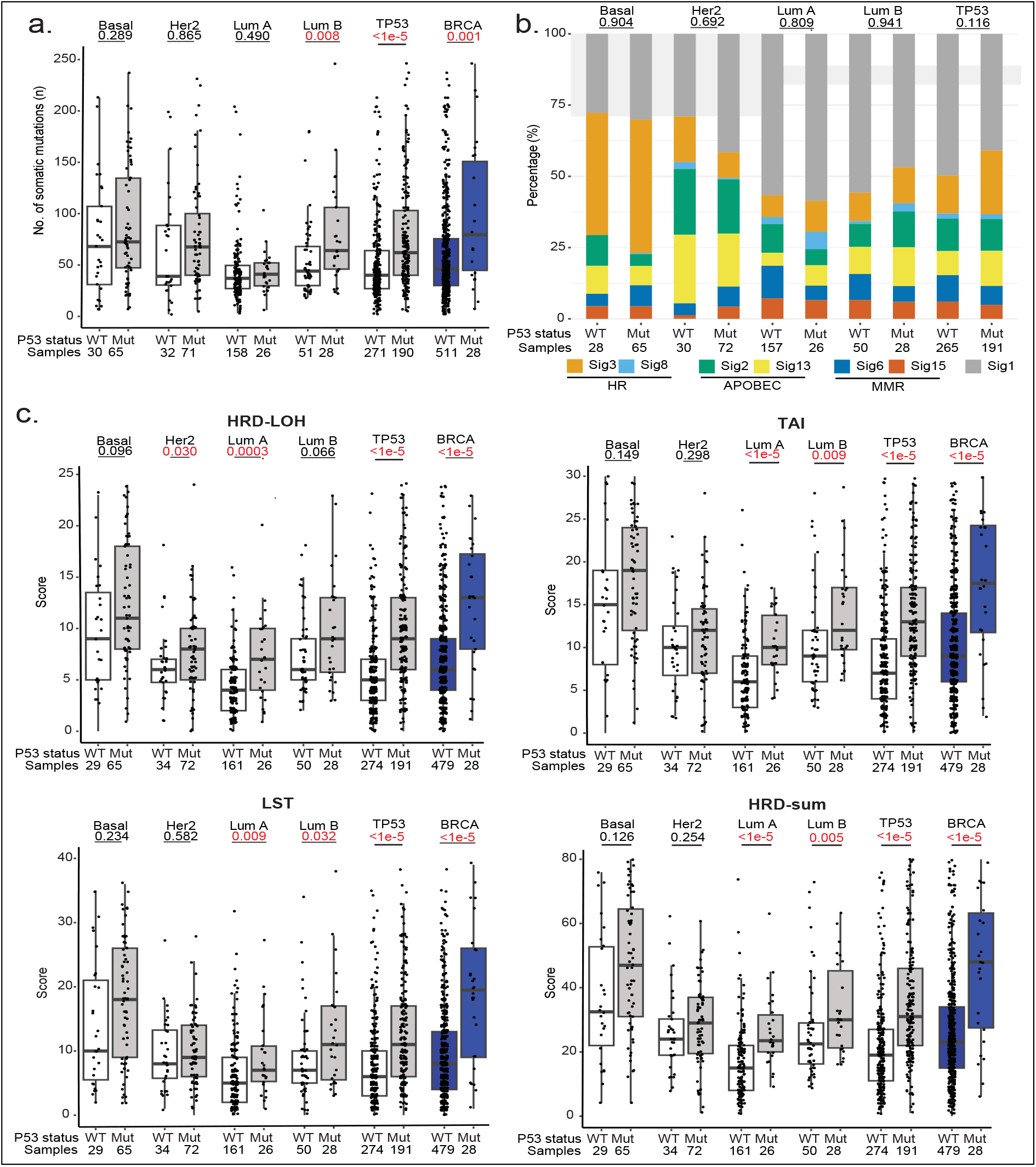
Comparison of mutational profiles for tumours with and without *TP53* somatic mutations. **a.** Total number of somatic alterations (single nucleotide variants (SNVs)) and indels identified in mutant and wild type *TP53* tumours, within each individual PAM50 subtype as well as overall (“TP53”), with germline *BRCA* mutated tumours (“BRCA”) included for comparison. **b.** The stacked bar plot demonstrates the proportion of major mutational signatures in between mutant *TP53* and wild type TP53 across subtypes. **c.** Comparison of genomic scar scores for mutant *TP53* tumours and wild type *TP53* tumours. Boxplots represent medians (centre line) and interquartile range, and whiskers represent the maximum and minimum values within 1.5 times the interquartile range from the edge of the box. Each data point represents an individual sample. P-values are derived from Mann–Whitney U tests for figures (a) and (c) and chi-square tests for figure (b).

Following that, we determined the proportion of the major mutational signatures in the tumour samples (Figure 1b). Mutational signatures are characteristic patterns of mutations associated with different types of DNA damage linked to various exogenous and endogenous mutagens, as well as DNA repair or replicative mechanisms (Helleday et al., 2014). These signatures, as defined by the COSMIC database (https://cancer.sanger.ac.uk/signatures/), include the age-related signature 1, homologous recombination (HR) deficiency-related signatures 3 and 8, MMR-related signatures 6 and 15 and APOBEC enzyme-related signatures 2 and 13 (Figure 1b). However, none of these signatures were significantly different between mutant and wild type *TP53* tumours across the subtypes, suggesting that *TP53* mutations have little effect on the mutational signatures of tumours (Figure 1b). We did note small increases in the HR deficiency-related signatures 3 and 8 when *TP53* is mutated in all subtypes except Her2-enriched, but these increases were not statistically significant.

We next examined other features of HR deficiency including genomic loss of heterozygosity (LOH), telomeric allelic imbalance (TAI) and large-scale state transition (LST), as well as an overall HRD-sum score that is the total of these scar signature scores (Abkevich et al., 2012;Birkbak et al., 2012; Popova et al., 2012; Telli et al., 2016). Inevitably, when aggregating across all subtypes, tumours with germline *BRCA* mutations and tumours with *TP53* somatic mutations had significantly higher HR deficiency scores compared to wild type tumours (*p* < 1e-5, Figure 1c). Nevertheless, when examined within subtypes, the increase in scores was more pronounced in luminal A and luminal B tumours, and less pronounced in basal-like and Her2-enriched tumours (Figure 1c). This is reflected in the overall HRD scores (HRD-sum), where, in subtype-specific comparisons, this measure was significantly different between mutant and wild type *TP53* only in luminal A and luminal B tumours (p < 1e-5 and p = 0.005), and not in the basal-like (p = 0.126) and Her2-enriched (p = 0.254) subtypes (Figure 1c).

Overall, these results suggest that *TP53* somatic mutations are associated with higher numbers of somatic mutations and higher HR-deficiency scores, as detectable by whole-exome sequencing. However, these associations appear to be stronger and more consistent in luminal A and B tumours and weaker in Her2-enriched and basal-like tumours.

### Differential Gene Expression Analysis

Next, to determine subtype-specific transcriptomic changes associated with of *TP53* somatic mutations, we employed RNA-Seq data to conduct differential gene expression analyses between tumours with and without *TP53* somatic mutations in each of the four main PAM50 molecular subtypes. The criteria for selecting differentially expressed genes (DEGs) were set as follows: i) absolute log_2_ fold change more than 1.5; ii) p-value less than 0.05. A comparison of upregulated and downregulated DEGs across subtypes revealed that there was remarkably little overlap between the different subtypes, with no genes that overlapped across all four molecular subtypes even when the high p-value threshold of 0.05 was used. Additionally, transcriptomic differences associated with *TP53* somatic mutations appeared to be particularly pronounced in the luminal A subtype, which had a much higher number of significantly downregulated genes compared to the other subtypes (Figure 2).

**Figure 2.**
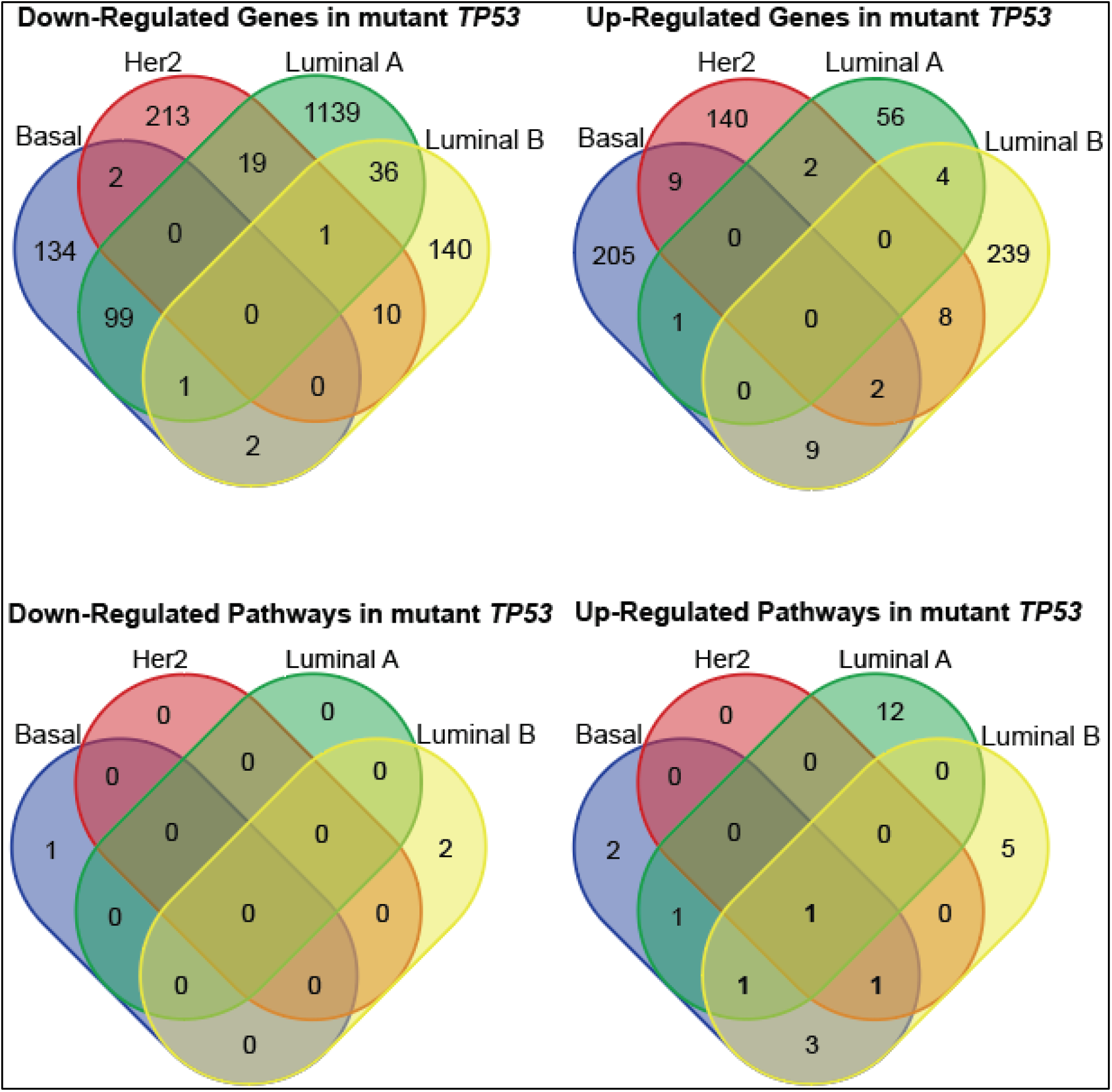
Differentially expressed genes and Hallmark pathways across Basal, Her2, Luminal A and Luminal B subtypes. (Top) The Venn diagrams show downregulated (left) and upregulated (right) genes across the Basal, Her2, Luminal A and Luminal B subtypes. The numbers indicate the number of overlapping differentially expressed genes across subtypes. (Bottom) The lower Venn diagrams display downregulated (left) and upregulated (right) Hallmark gene sets according to gene set enrichment analysis (GSEA) across subtypes. Only genes and Hallmark gene sets with p-value < 0.05 were included.

To understand this further, we conducted pathway analyses using gene set enrichment analysis (GSEA) (Mootha et al., 2003; Subramanian et al., 2005). To investigate the overall biological impact of *TP53* somatic mutations on the tumours of MyBrCa patients, we focused on the Hallmark gene sets curated by MSigDB (Liberzon et al., 2015). Comparison of the downregulated and upregulated pathways in tumours with mutant *TP53* revealed only one pathway that was consistently upregulated across all subtypes (mTORC1 Signaling), and two pathways that were upregulated in three subtypes (Glycolysis and UV Response Up). Overall, there were 15 Hallmark pathways with significant differences in expression between mutant and wild type *TP53* in luminal A, 13 in luminal B, 10 in basal-like, and only 2 in Her2-enriched. This analysis suggests that the effect of *TP53* mutations is subtype-specific, as there were very few overlapping gene sets across molecular subtypes, and the subtypes are observed to have largely distinct upregulated or downregulated pathways when *TP53* is mutated (Figure 2).

Together, these results suggest that *TP53* somatic mutations are associated with changes to the tumour transcriptome that vary by breast cancer subtype, with surprisingly little overlap. The DEG and GSEA results also suggest that the association between *TP53* somatic mutations and pathway dysregulation is most pronounced in the luminal A subtype, and least pronounced in the Her2-enriched subtype.

### Single-sample Pathway Analysis

To validate and further explore the important pathways associated with mutant *TP53* across subtypes, we performed single-sample GSEA (ssGSEA) analyses. We analyzed the ssGSEA results for the hallmark pathways that were observed to overlap across three or four subtypes (mTORC1 Signaling, Glycolysis and UV Response Up pathways). We also examined three other Hallmark pathways (P53 Pathway, DNA Repair, and G2M Checkpoint) that are associated with known roles of the *TP53* gene such as DNA repair and cell cycle.

The ssGSEA analyses showed that *TP53* somatic mutations were consistently associated with the mTORC1 Signaling and Glycolysis pathways across all subtypes, but found that the UV Response Up pathway was significantly different only in the luminal A subtype (Figure 3). Additionally, our analyses also showed that the P53 pathway was not significantly dysregulated when comparing mutant and wild type *TP53* samples within each subtype, while the DNA Repair and G2M Checkpoint pathways had inconsistent associations with *TP53* somatic mutations (DNA Repair: significantly upregulated only in luminal B (p = 0.027), G2M Checkpoint: significantly upregulated in luminal A (p = 0.012) and basal-like (p = 0.042); Figure 3). Notably, aggregating the data for mutant versus wild type *TP53* across all subtypes often led to different results in terms of statistical significance and/or the direction of the effect due to the higher prevalence of *TP53* somatic mutations in the basal and Her2-enriched subtypes coupled with differences in gene set expression between the subtypes, emphasizing the importance of including tumour subtype as a covariate.

**Figure 3.**
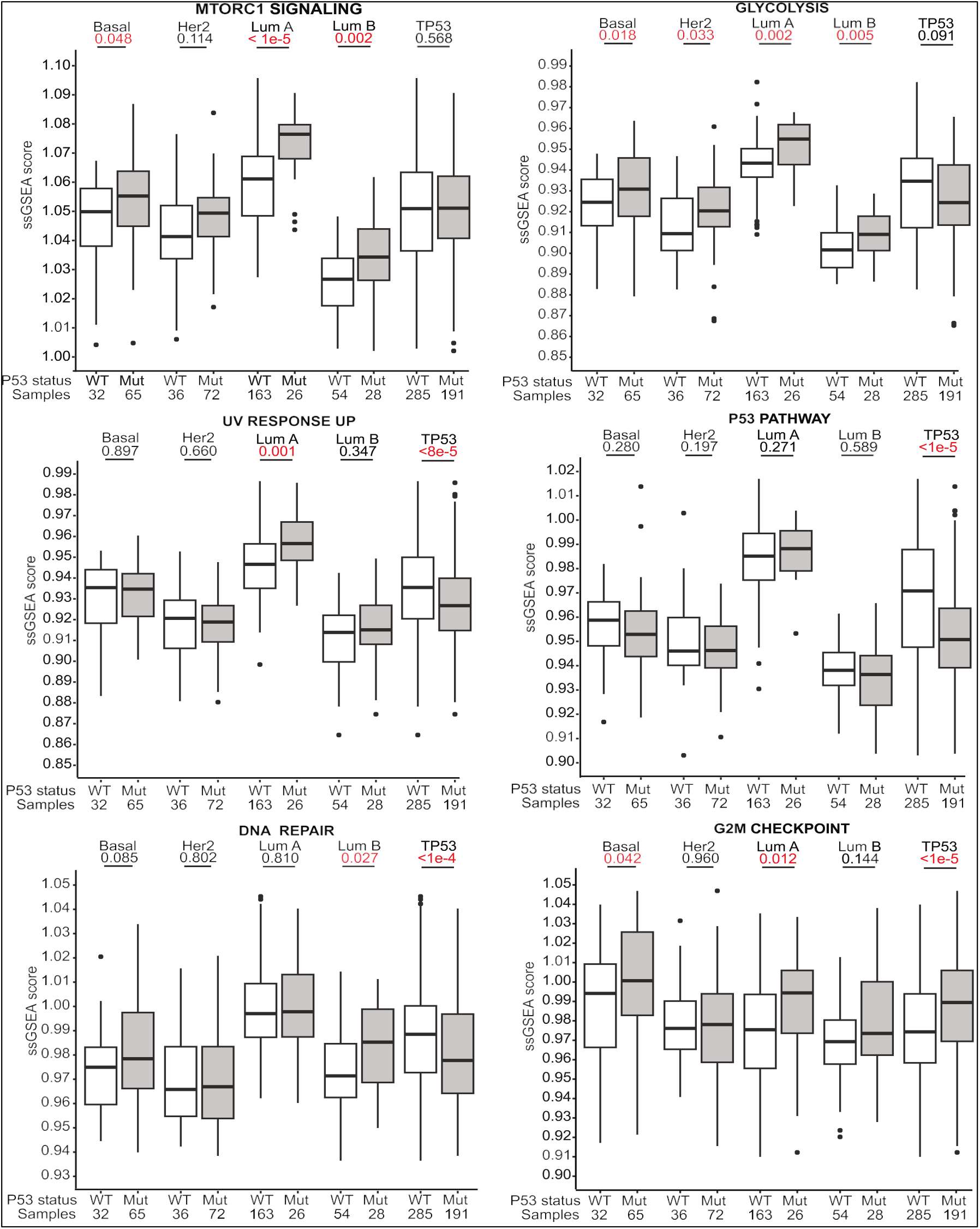
Pathway expression in tumours with and without *TP53* somatic mutations in different breast cancer subtypes. Boxplots compare single-sample GSEA scores for various Hallmark pathways in mutant and wild type *TP53* tumours across different PAM50 subtypes as well as overall (“TP53”). Hallmark pathways were selected if they were previously found to be dysregulated across three or more subtypes in GSEA analysis, or based on known functions of the *TP53* gene. Boxplots represent medians (centre line) and interquartile range, and whiskers represent the maximum and minimum values within 1.5 times the interquartile range from the edge of the box. Each data point represents an individual sample. P-values are derived from Mann–Whitney U tests.

In summary, these data suggest that, across all molecular subtypes, *TP53* somatic mutations are most consistently associated with upregulation of the mTORC1 signaling pathway, as well as the glycolysis pathway to a lesser extent. On the other hand, *TP53* somatic mutations have only small and inconsistent associations with transcriptional alterations in the p53, DNA repair and cell cycle pathways. Similar to our previous results, the differences in pathway expression between samples with mutant and wild type *TP53* appear to be most pronounced in the luminal A subtype and least pronounced in the Her2-enriched subtype.

### Cancer Cell Fraction

Next, we hypothesized that subtype-specific differences in the mutational and transcriptional profile of tumours with *TP53* somatic mutations may be due to *TP53* somatic mutations being driver versus passenger mutations in the different molecular subtypes. To assess this hypothesis, we analyzed data on the mutational cancer cell fraction (CCF) of *TP53* in individual tumour samples that was generated by Pan et al. (2020) from copy number data using an ASCAT pipeline. Interestingly, a comparison of *TP53* CCF across breast cancer subtypes showed that the Her2-enriched and basal-like subtypes had lower *TP53* CCFs compared to luminal A and B (Supplementary Figure 2). This finding is consistent with *TP53* mutations acting as driver mutations in luminal A and luminal B tumours but as passenger mutations in the basal-like and Her2-enriched subtypes.

## Discussion

In this study, we compared the molecular profiles of tumours with and without *TP53* somatic mutations in a cohort of 489 Malaysian breast cancer patients. Using whole exome and transcriptome data, we conducted an analysis of tumour mutational burden, mutational signatures, HR deficiency scores, and differentially expressed genes and pathways. Our analyses resulted in two main findings: i) the association between *TP53* somatic mutations and HR deficiency scores is stronger in the luminal A and luminal B subtypes, ii) *TP53* mutations are associated with subtype-specific gene expression differences that are more pronounced in the luminal A and luminal B subtypes, with only the mTORC1 and glycolysis pathways being consistently dysregulated across most subtypes when *TP53* is mutated. We suggest that these results may be due to due to *TP53* somatic mutations generally being driver mutations in luminal A and luminal B tumours as opposed to sometimes being passenger mutations in the basal-like and Her2-enriched subtypes. Indeed, the cancer cell fraction analyses suggest that *TP53* mutations may act as driver mutations in luminal A and luminal B tumours but as passenger mutations in the basal-like and Her2-enriched subtypes. Taken together, these results highlight the importance of considering molecular subtype when examining the role of *TP53* in breast cancer. These results also suggest that therapies for Asian breast cancer that target p53 or other downstream pathways may be more effective in the luminal A and luminal B tumour subtypes.

Given that TP53 is known to have important roles in DNA repair and the maintenance of genomic stability (Lane and Levine, 2010; Silwal-Pandit et al., 2017), our finding that *TP53* somatic mutations are associated with an increase in tumour mutation burden in some subtypes is not surprising. Similarly, previous research has also shown that *TP53* mutations are associated with chromosomal instability and high HRD scores (Donehower et al., 2019;Knijnenburg et al., 2018). Our results confirm that these associations are also present in a large cohort of Asian breast tumour samples.

Our results also indicate a strong association between *TP53* somatic mutations and the mTORC1 signaling and glycolysis pathways in Asian breast cancer. The p53 protein is well known to inhibit mTORC1 signaling through multiple mechanisms (Feng et al., 2005; Gwinn et al., 2008; Hasty et al., 2013) in response to cellular stresses such as DNA damage. Similarly, *TP53* mutations have been shown to affect energy metabolism at multiple levels in TCGA breast cancer samples and mutant breast cancer cell lines (Harami-Papp et al., 2016; Liu et al., 2015).

On the other hand, the weak association between *TP53* somatic mutation and p53 signaling, DNA repair, and cell cycle pathways is surprising, given the known functions of TP53. The lack of association between *TP53* mutations and p53 signaling has been noted before in other cohorts (Leroy et al., 2014), and may be due to the existence of compensatory mechanisms (Soussi and Kroemer, 2018; Wasylishen and Lozano, 2016). However, other studies have found strong associations between *TP53* somatic mutations and transcriptional changes affecting cell cycle progression (Donehower et al., 2019). Further research will be necessary to determine if our results can be generalized to the wider Asian population. Notably, when the data for mutant versus wild type *TP53* is aggregated across all subtypes, the results often lead to misleading conclusions due to differences in sample size, prevalence of *TP53* mutations and expression of various gene sets between the various subtypes, highlighting the importance of doing these comparisons in a subtype-specific manner.

Perhaps the most surprising aspect of our results is that the associations between *TP53* somatic mutations and genomic and transcriptomic changes are strongest in the luminal A and B subtypes and weaker in the basal-like and Her2-enriched subtypes. This is surprising because the prevalence of *TP53* somatic mutations is generally lower in the luminal A and B subtypes and higher in the basal-like and Her2-enriched subtypes, such that *TP53* has been considered to be a less important driver gene for luminal A and luminal B subtypes (Silwal-Pandit et al.,2017). Our results, on the other hand, suggest that this paradigm may not be as applicable to luminal A and luminal B breast cancers arising in the Asian population, where *TP53* may have a stronger driver role. This may be related to the finding by previous studies that ER+ tumours from Asian breast cancer patients have higher frequencies of *TP53* somatic mutations compared to ER+ tumours from Western studies, although the mechanism behind this population-specific effect, whether genetic or environmental, remains to be elucidated (Kan et al., 2018; Pan et al., 2020). Other possible explanations for the lower strength of associations in basal-like and Her2-enriched subtypes are that basal-like tumours are known to have high levels of intra-tumoural heterogeneity, thus diluting the transcriptional effects of sub-clonal *TP53* mutations, while Her2-enriched tumours are primarily driven by *ERBB2* copy number amplification events which renders *TP53* mutations less important.

In conclusion, *TP53* somatic mutations in Asian breast cancer are associated with HR deficiency and upregulation of the mTORC1 signaling and glycolysis pathways. These associations appear to be stronger in luminal A and luminal B tumours, and weakest in the Her2-enriched subtype, which may be an important consideration for therapies that target *TP53* or other downstream pathways in the Asian population. These results may also provide useful context for further research into *TP53* somatic mutations as predictive or prognostic biomarkers for breast cancers arising in the Asian population.

## Materials and Methods

### Whole exome and transcriptome dataset from the MyBrCa cohort

Our study employed whole exome and transcriptome data of tumour samples collected from the MyBrCa cohort that has been described previously (Pan et al., 2020). The MyBrCa tumour genomics cohort comprised of 560 breast cancer patients recruited as part of the Malaysian Breast Cancer (MyBrCa) study (Tan et al., 2018) at the Subang Jaya Medical Centre, whose fresh frozen primary tumours were selected to undergo deep whole exome and transcriptome sequencing. Genomics analyses of these patient tumour samples have been approved by the Ethics Committee of Subang Jaya Medical Centre (reference no: 201208.1). More specific details on the sequencing methodology are available from the original publication.

Among the 560 patients, 29 tumour samples that had *TP53* mutations with unknown or uncertain significance or conflicts over its pathogenicity (VUS) were excluded from this study. Only pathogenic/likely pathogenic somatic mutations were included in this study. Another 39 patients without available WES or RNAseq data were excluded from further analysis for a final sample size of 492 samples. No overlapping sample was included in each set. Tumour samples were categorised as either mutant or wild type *TP53*.

### Mapping and variant calling of *TP53* mutations

Analysis of sequencing data was performed as described previously in study by Pan et al. (2020). In summary, for WES, sequenced reads were aligned to the human genome b37 plus decoy genome by utilising bwa-mem version 0.7.12 (Pan et al., 2020). Local realignment, duplicate removal and base quality recalibration were performed via the Genome Analysis Toolkit (GATK, v3.1.1) (McKenna et al., 2010). Somatic SNVs were identified via GATK3 Mutect2 (McKenna et al., 2010), whereas small insertions and deletions (indels) were established by Strelka2 (Kim et al., 2018).

The variants present only in tumour tissue samples were consequently categorised as somatic mutations. The reference sequences for numbering were based on the NCBI GenBank Database for TP53 (NM_000546.6). Clinical variant annotations were obtained from NCBI ClinVar (http://www.ncbi.nlm.nih.gov/clinvar) and Varsome (https://varsome.com/). The variants are considered as pathogenic mutations if they were annotated as “pathogenic” in NCBI ClinVar.

### Mutational signatures

Mutational signatures in each breast tumour sample were determined from annotated VCF files using deconstructSigs (Rosenthal et al., 2016), using the single base-pair substitution (SBS) signatures described in the COSMIC database.

### HR deficiency scores

The following measures of HR deficiency were established as depicted earlier: (1) LOH, (2) LST and, (3) TAI (Birkbak et al., 2012; Telli et al., 2016). Allele-specific copy number (ASCN) profiles on paired normal-tumour BAM files were classified via Sequenza (Favero et al., 2015) and utilised to analyse the single measure scores and HRD-sum scores via the scarHRD R package (Sztupinszki et al., 2018).

### Gene Expression Analysis

Gene expression data used for this study was the same as in the original analysis (Pan et al. 2020). Briefly, RNA-seq reads were mapped to the hs37d5 human genome and the ENSEMBLE GrCh37 release 87 human transcriptome via the STAR aligner (v.2.5.3a) (Dobin et al., 2013). Variant calling for RNA-seq data was also performed by utilising using the GATK Best Practices workflow for RNA-seq. Gene-level transcript counts were obtained using featureCounts (v. 1.5.3).

### Molecular classification based on gene expression data

Gene-level count matrices for the cohort were transformed into log2 counts-per-million (logCPM) using the voom function from the limma (v. 3.34.9) R package. The transformed matrices were then was subtyped according to PAM50 designations using the Genefu package in R (v. 2.14.0).

### Differential gene expression and functional enrichment analysis

Gene expression was analysed with the DEseq2 package, an R-based open-source software designed to analyse transcriptomic data for differential expression, as previously described (Love et al., 2014). GSEA was then performed for each downregulated and upregulated genes from each PAM50 subtypes using Hallmark Gene set (Liberzon et al., 2015; Mootha et al.,2003; Subramanian et al., 2005). Enrichment analysis was done with default parameter settings. An enrichment score was calculated for each gene set (KS-statistics) reflecting if the genes in the particular gene set appeared in the top (positive score), in the bottom (negative score), or were randomly distributed (close to zero score). These scores were compared with scores calculated from 1,000 randomly permuted gene lists, in order to calculate false discovery rates (FDR) (cutoff at FDR=0.05). The single sample gene set enrichment analysis (ssGSEA) was applied to analyze the RNA-seq data as well. Hallmark gene sets from the Molecular Signatures Database were used for GSEA and ssGSEA analysis (Liberzon et al., 2015; Mootha et al., 2003; Subramanian et al., 2005).

### Statistical analysis

The Mann–Whitney *U* test and the Chi-square test were executed for comparisons of variables between categories. *P* < 0.05 was considered statistically significant and all tests were two-sided. Statistical analyses were performed using R v4.0. All box and whiskers plots in the main and supplemental figures are constructed with boxes indicating 25th percentile, median and 75th percentile, and whiskers showing the maximum and minimum values within 1.5 times the inter-quartile range from the edge of the box, with outliers not shown.

## Data and Code Availability

The data that support the findings of this study are available on the European Genome-phenome Archive under the study accession number EGAS00001004518. Access to controlled patient data will require the approval of the Data Access Committee. Further details and other data that support the findings of this study are available from the lead contact upon request.

## Acknowledgements

Cancer Research Malaysia receives charitable funding from the Scientex Foundation, Estée Lauder Companies, Yayasan Petronas, and Yayasan Sime Darby, which contributed to the funding of this study. The authors would like to thank Dr. Muhammad Mamduh Ahmad Zabidi, Mei-Yee Meng, and Siti Norhidayu Hasan, and the Subang Jaya Medical Centre Tissue Diagnostics laboratory for contributing to the initial MyBrCa tumour sequencing dataset used in this study. We would also like to acknowledge the contribution of Bethan Sandey, Oscar Rueda, Suet-Feung Chin, and Carlos Caldas from the Caldas Lab, along with the Core Genomics Facility at the CRUK Cambridge Institute, towards the sequencing of the samples.

## Author Contributions

M.E.R. performed all the data analysis and wrote the manuscript. J.M.C.L contributed to data analysis. J.M.C.L, P.S.N., C.H.Y. and P.R. contributed to sample collection and processing and data collection. S.H.T. and J.W.P. designed experiments, interpreted results, guided the data analysis, and drafted the manuscript. The project was directed and co-supervised by S.H.T. and J.W.P., and were responsible for final editing.

