## Supplementary figures and images for "*TP53* somatic mutations in Asian breast cancer are associated with subtype-specific effects"

### Supplemental Figure 1

Supplementary Figure 1. Distribution of somatic TP53 mutations identified in MyBrCa Cohort

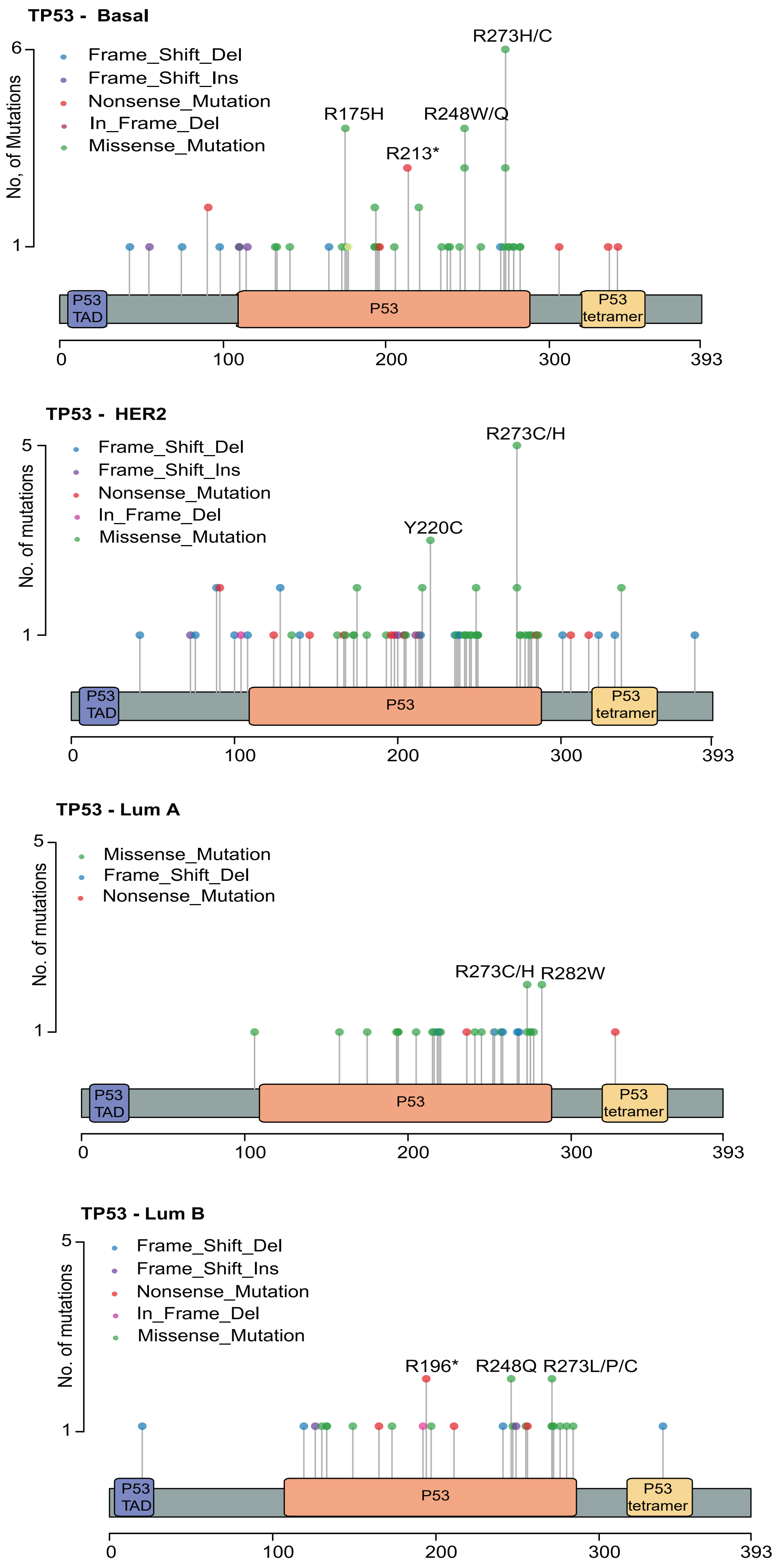

### Supplemental Figure 2

**Supplementary Figure 2. *TP53* Cancer Cell Fraction**

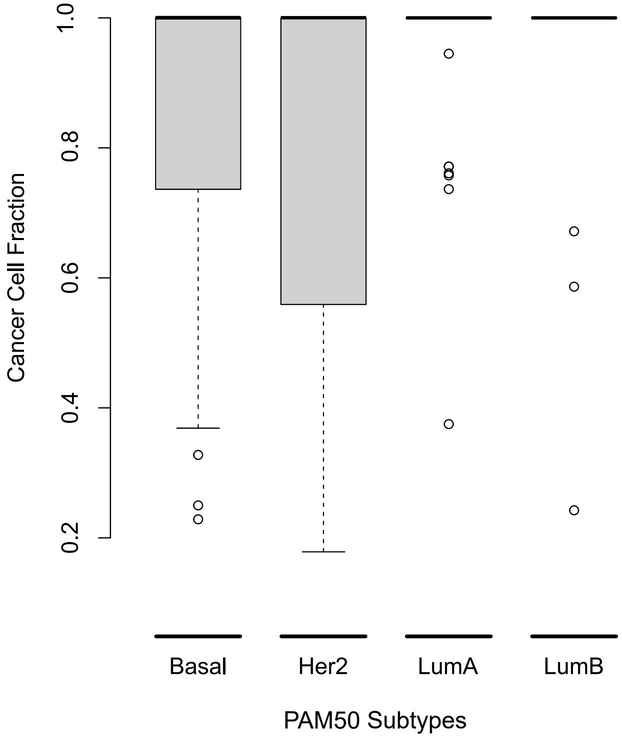
