## Supplemental Table 1 for "*TP53* somatic mutations in Asian breast cancer are associated with subtype-specific effects"

**Supplementary Table 1.** Clinically relevant or potentially clinically relevant somatic mutations of by subtype

| cDNA_Change | Protein_Change | Prevalence (%) | | | | Statistical Significance |
|---|---|---|---|---|---|---|
| | | Basal (n=65) | Her2 (n=72) | LumA (n=26) | LumB (n=28) | |
| **c.524G>A** | p.R175H | 6.20 | 2.80 | 3.85 | 3.57 | 0.797 |
| **c.586C>T** | p.R196* | 1.50 | 1.39 | 0.00 | 7.69 | 0.230 |
| **c.614A>G** | p.Y205C | 1.50 | 1.39 | 3.85 | 0.00 | 0.721 |
| **c.637C>T** | p.R213* | 4.62 | 1.39 | 0.00 | 3.57 | 0.524 |
| **c.659A>G** | p.Y220C | 3.07 | 4.17 | 3.85 | 0.00 | 0.753 |
| **c.743G>A** | p.R248Q | 6.15 | 2.78 | 0.00 | 7.14 | 0.435 |
| **c.817C>T** | p.R273C | 4.62 | 6.95 | 7.69 | 3.57 | 0.857 |
| **c.818G>A** | p.R273H | 9.23 | 2.78 | 3.85 | 0.00 | 0.173 |

Above table shows the top ten most common mutations across pam50 subtypes. The 273C and R175H mutations can be observed across all the subtypes and known to be common hotspot mutation in TP53.
