## Supplemental Table 2 for "*TP53* somatic mutations in Asian breast cancer are associated with subtype-specific effects"

**Supplementary Table 2** GSEA Hallmark Pathway

| Subtype | Regulation | Hallmark Pathway | NES | FDR | p-val |
|---|---|---|---|---|---|
| Basal | Up-regulated | HALLMARK_UNFOLDED_PROTEIN_RESPONSE | 1.863 | 0.051 | 0.004 |
| Basal | Up-regulated | HALLMARK_MYC_TARGETS_V2 | 1.768 | 0.041 | 0.006 |
| Basal | Up-regulated | HALLMARK_MYC_TARGETS_V1 | 1.847 | 0.020 | 0.006 |
| Basal | Up-regulated | HALLMARK_E2F_TARGETS | 1.860 | 0.026 | 0.010 |
| Basal | Up-regulated | HALLMARK_UV_RESPONSE_UP | 1.570 | 0.104 | 0.016 |
| Basal | Up-regulated | HALLMARK_GLYCOLYSIS | 1.682 | 0.052 | 0.016 |
| Basal | Up-regulated | HALLMARK_MTORC1_SIGNALING | 1.713 | 0.044 | 0.025 |
| Basal | Up-regulated | HALLMARK_HYPOXIA | 1.546 | 0.106 | 0.027 |
| Basal | Up-regulated | HALLMARK_G2M_CHECKPOINT | 1.724 | 0.048 | 0.030 |
| Basal | Down-regulated | HALLMARK_PANCREAS_BETA_CELLS | -1.458 | 1.000 | 0.046 |
| Her2 | Up-regulated | HALLMARK_MTORC1_SIGNALING | 1.680 | 0.507 | 0.011 |
| Her2 | Up-regulated | HALLMARK_GLYCOLYSIS | 1.657 | 0.312 | 0.013 |
| Luminal A | Up-regulated | HALLMARK_CHOLESTEROL_HOMEOSTASIS | 1.713 | 0.082 | 0.003 |
| Luminal A | Up-regulated | HALLMARK_IL2_STAT5_SIGNALING | 1.693 | 0.080 | 0.003 |
| Luminal A | Up-regulated | HALLMARK_INFLAMMATORY_RESPONSE | 1.817 | 0.142 | 0.005 |
| Luminal A | Up-regulated | HALLMARK_COMPLEMENT | 1.805 | 0.079 | 0.005 |
| Luminal A | Up-regulated | HALLMARK_INTERFERON_GAMMA_RESPONSE | 1.803 | 0.053 | 0.014 |
| Luminal A | Up-regulated | HALLMARK_KRAS_SIGNALING_UP | 1.677 | 0.068 | 0.014 |
| Luminal A | Up-regulated | HALLMARK_PI3K_AKT_MTOR_SIGNALING | 1.582 | 0.089 | 0.018 |
| Luminal A | Up-regulated | HALLMARK_IL6_JAK_STAT3_SIGNALING | 1.674 | 0.062 | 0.024 |
| Luminal A | Up-regulated | HALLMARK_MTORC1_SIGNALING | 1.639 | 0.077 | 0.028 |
| Luminal A | Up-regulated | HALLMARK_TNFA_SIGNALING_VIA_NFKB | 1.595 | 0.088 | 0.030 |
| Luminal A | Up-regulated | HALLMARK_HYPOXIA | 1.419 | 0.154 | 0.033 |
| Luminal A | Up-regulated | HALLMARK_ALLOGRAFT_REJECTION | 1.733 | 0.087 | 0.033 |
| Luminal A | Up-regulated | HALLMARK_UV_RESPONSE_UP | 1.462 | 0.138 | 0.037 |
| Luminal A | Up-regulated | HALLMARK_EPITHELIAL_MESENCHYMAL_TRANSITION | 1.692 | 0.068 | 0.038 |
| Luminal A | Up-regulated | HALLMARK_APICAL_JUNCTION | 1.489 | 0.126 | 0.050 |
| Luminal B | Up-regulated | HALLMARK_OXIDATIVE_PHOSPHORYLATION | 2.012 | 0.007 | 0.000 |
| Luminal B | Up-regulated | HALLMARK_MYC_TARGETS_V1 | 1.949 | 0.006 | 0.000 |
| Luminal B | Up-regulated | HALLMARK_MYC_TARGETS_V2 | 1.913 | 0.008 | 0.000 |
| Luminal B | Up-regulated | HALLMARK_FATTY_ACID_METABOLISM | 1.794 | 0.024 | 0.000 |
| Luminal B | Up-regulated | HALLMARK_MTORC1_SIGNALING | 1.896 | 0.007 | 0.004 |
| Luminal B | Up-regulated | HALLMARK_UNFOLDED_PROTEIN_RESPONSE | 1.742 | 0.036 | 0.011 |
| Luminal B | Up-regulated | HALLMARK_GLYCOLYSIS | 1.543 | 0.097 | 0.022 |
| Luminal B | Up-regulated | HALLMARK_ADIPOGENESIS | 1.622 | 0.062 | 0.025 |
| Luminal B | Up-regulated | HALLMARK_DNA_REPAIR | 1.657 | 0.060 | 0.032 |
| Luminal B | Up-regulated | HALLMARK_UV_RESPONSE_UP | 1.457 | 0.133 | 0.042 |
| Luminal B | Up-regulated | HALLMARK_PEROXISOME | 1.417 | 0.149 | 0.049 |
| Luminal B | Down-regulated | HALLMARK_NOTCH_SIGNALING | -1.843 | 0.043 | 0.004 |
| Luminal B | Down-regulated | HALLMARK_WNT_BETA_CATENIN_SIGNALING | -1.587 | 0.302 | 0.039 |
